# Extending the Gaussian Network Model: Integrating Local, Allosteric, and Structural Factors for Improved Residue-Residue Correlation Analysis

**DOI:** 10.1101/2025.05.12.653468

**Authors:** Burak Erman

## Abstract

The Gaussian Network Model (GNM) has been successful in explaining protein dynamics by modeling proteins as elastic networks of alpha carbons connected by harmonic springs. However, its uniform interaction assumption and neglect of higher-order correlations limit its accuracy in predicting experimental B-factors and residue cross-correlations critical for understanding allostery and information transfer. This study introduces an information-theoretic enhancement to the GNM, incorporating mutual information-based corrections to the Kirchhoff matrix to account for multi-body interactions and contextual residue dynamics. By iteratively optimizing B-factor predictions and applying a Monte Carlo-driven maximum entropy approach to refine covariances, our method achieves significant improvements, reducing RMSDs between predicted and experimental B-factors by 26–46% across eight representative proteins. The model contextualizes residue assignments based on local density, secondary structure, solvent exposure, and allosteric roles, showing complex dynamic patterns beyond simple neighbor counts. Enhanced predictions of mutual information and entropy transfer in proteins like KRAS highlight improved capture of allosteric communication pathways. This evolvable framework, capable of incorporating additional effects and utilizing contextual residue assignments, enables precise studies of mutation effects on protein dynamics, with improved cross-correlation predictions potentially increasing accuracy in drug design and function prediction.

## 1. Introduction

The Gaussian Network Model (GNM)[1] emerged from the theoretical framework of polymer physics, specifically drawing upon the Constrained Junction Theory of rubber elasticity pioneered by Flory[2] and later refined in the Flory-Erman Constrained Junction Model (CJM)[3, 4]. This intellectual lineage— transferring concepts from synthetic polymer networks to biological macromolecules—provided crucial insights into the relationship between molecular topology and conformational dynamics. By conceptualizing folded proteins as elastic networks with alpha carbons serving as junction points connected by fictitious springs, the GNM successfully captured the essence of protein fluctuations governed by local crowding effects.

While the original formulation of the GNM has proven effective in predicting experimental B-factors using just a single adjustable parameter, it showed the possibility of rapidly predicting cross-correlations of fluctuations through the inversion of the Kirchoff matrix—a critical advancement for understanding allostery, information transfer pathways, and protein function more generally. This predictive capability has been particularly valuable, as direct experimental observation of such cross-correlations is only now emerging, with available data still relatively scarce[5, 6].

Systematic deviations in loop regions and densely packed structural elements have been observed when comparing GNM predictions with experimental B-factors, pointing to limitations in the current formulation of the Kirchhoff matrix. Addressing these discrepancies is critical for obtaining more accurate B-factor predictions, which would consequently lead to more reliable cross-correlation estimations that better capture true protein dynamics. While molecular dynamics simulations can provide such detailed correlations, GNM offers a significantly faster computational approach to obtaining this essential functional information.

This paper presents an information-theoretic approach to enhancing the GNM through iterative correction of the Laplacian elements, explicitly accounting for higher-order correlations between residues while preserving the computational efficiency of the Gaussian framework. Once a near-optimal B-factor profile is achieved, a Monte Carlo algorithm refines the cross-correlations by maximizing entropy while keeping the optimized B-factors fixed. This entropy maximization ensures the least biased Kirchhoff matrix, minimizing unsupported assumptions such as uniform interaction strengths or neglect of many-body correlations. The approach provides a straightforward method to compute off-diagonal covariance terms—critical for understanding protein dynamics—where their absence from experimental data underscores the importance of accurate computational evaluation.

An elastic polymeric network consists of chains linked to each other by covalent junctions. Alternatively, such a network may be conceptualized as a single long primary chain folded in three-dimensional space and covalently connected to itself at distinct points forming junctions. Each junction fluctuates within a coordination sphere defined by a specific cutoff radius, where approximately 50-100 other junctions may be present, many originating from topologically distant segments. Consequently, junction fluctuations are governed by both network connectivity and the degree of interpenetration within this coordination sphere[3, 7]. The mean-square fluctuation of each junction exhibits an inverse relationship with the total number of neighboring junctions, both covalently bonded and non-bonded, that interpenetrate the coordination sphere. The statistical mechanical framework of the CJM has demonstrated remarkable success in explaining virtually all observed behaviors of polymeric elastomers[8-13].

The GNM extends this paradigm to proteins, with alpha carbons assuming the role of junctions[1]. Each alpha carbon interacts with two types of neighbors: those covalently connected along the primary chain and those originating from topologically distant segments. Every alpha carbon occupies a well-defined equilibrium position around which it undergoes thermal fluctuations. Thus, analogous to elastic networks, proteins function as deformable elastic bodies whose elasticity is predominantly determined by these fluctuations. Following the CJM approach, the GNM mathematically implements the physical principle that the mean squared fluctuation of an alpha carbon varies inversely with the total number of neighboring alpha carbons, both covalent and non-covalent. The validity of this physical principle is shown in Figure 1 results are shown for several examples.

**Figure 1.**
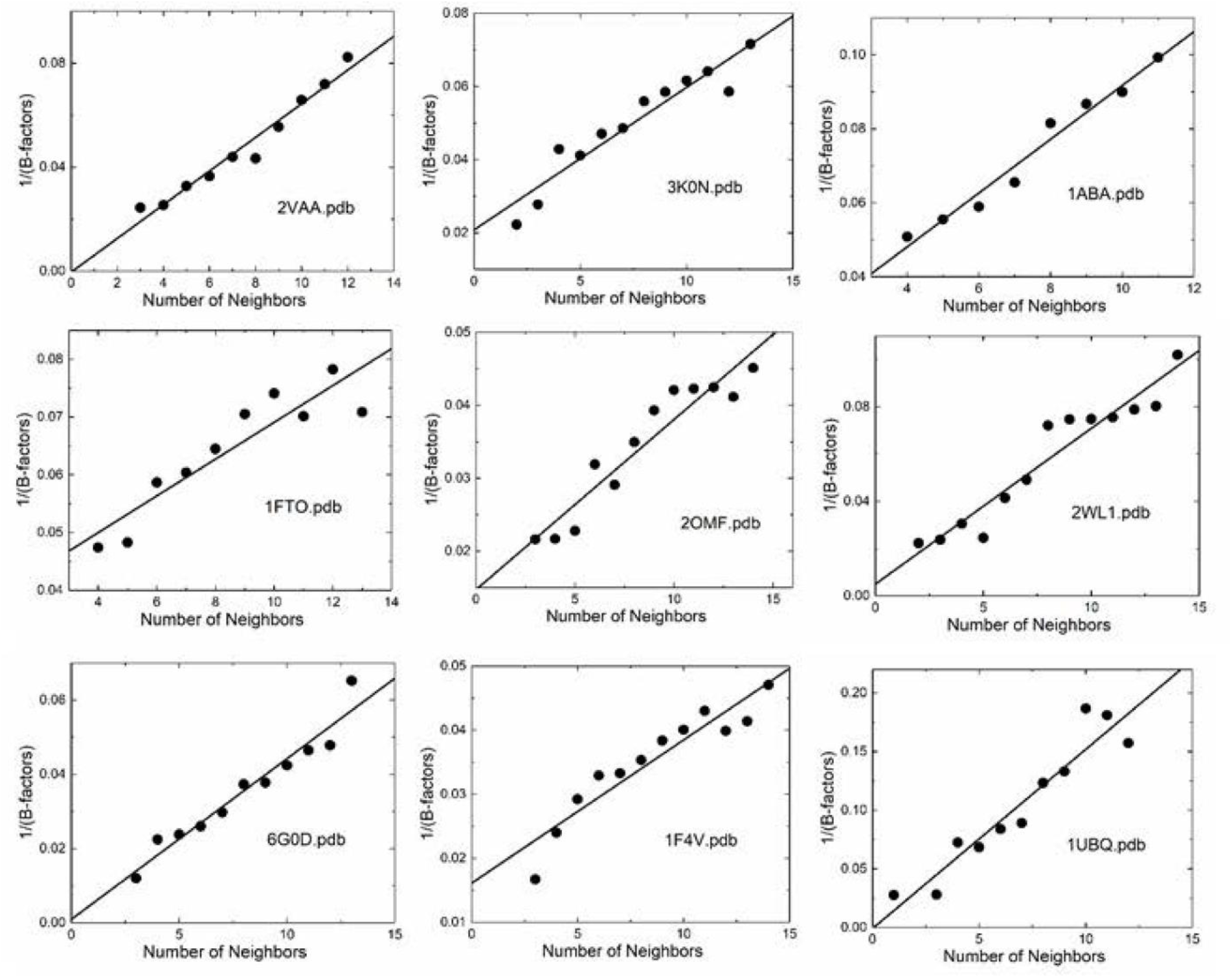
Inverse Relationship of B-Factors and Neighbor Counts. A scatter plot of neighbor counts (number of alpha carbons within a 7 Å radius) versus inverse B-factors (1/B-factor, Å^−2^). A least squares regression line is fitted to the data to model the relationship between neighbor counts and inverse B-factors, minimizing the sum of squared differences between the observed inverse B-factors and the predicted values from the linear model. The linear trend demonstrates that residues with fewer neighbors (sparse) exhibit higher B-factors, while residues with many neighbors (crowded) show lower B-factors. Each data point represents the mean B-factor for residues having a specific neighbor count.

This figure supports Flory’s hypothesis that residues in crowded regions fluctuate less, extending beyond elastic polymeric networks to describe coarse-grained protein behavior. The observed scatter in the figure reflects that while neighbor count provides a simple metric of local packing density, B-factors incorporate numerous additional contributions including, but not limited to, secondary structure elements, solvent accessibility, and nonlinear dynamic effects—complex factors that cannot be captured by neighbor counts alone.

This completes the analogy wherein a protein is represented as a coarse-grained polymer chain folded onto itself, with alpha carbons fluctuating about their respective equilibrium positions. Both CJM and GNM assume multivariate Gaussian distributions for these fluctuations, resulting in linear spring-like forces between pairs. Molecular dynamics simulations largely validate this assumption, demonstrating that protein fluctuations indeed approximate Gaussian behavior, consistent with expectations from the central limit theorem[14].

Paul Flory’s groundbreaking insight explained the inverse relationship between junction fluctuations and the degree of interpenetration from neighboring junctions in elastic networks. This fundamental observation, underappreciated at its inception[15], exemplifies Flory’s extraordinary scientific foresight. The principle has transcended its original context to become foundational in our understanding of protein dynamics through the GNM, establishing a crucial bridge between polymer physics and structural biology.

Figure 2 further illustrates the effect of crowding on fluctuations by comparing experimentally observed B-factor profiles (thin lines) with the inverse of alpha carbon neighbor counts for each residue (thick lines), though this simplified metric excludes the influence of network connectivity present in the full GNM. The thick lines, derived simply by calculating the reciprocal of the number of neighboring alpha carbons (degree of interpenetration) and least-squares fitted to experimental data, demonstrate the significant contribution of local crowding to residue fluctuations. Notably, these curves reflect only local neighbor effects; contributions from connectivity, represented by the inverse Kirchhoff matrix, are not incorporated. The GNM extends this local crowding effect by integrating the protein’s complete interaction network, generalizing the Kirchhoff matrix approach from phantom network theory [16, 17] to encompass both covalent and non-covalent interactions.

**Figure 2.**
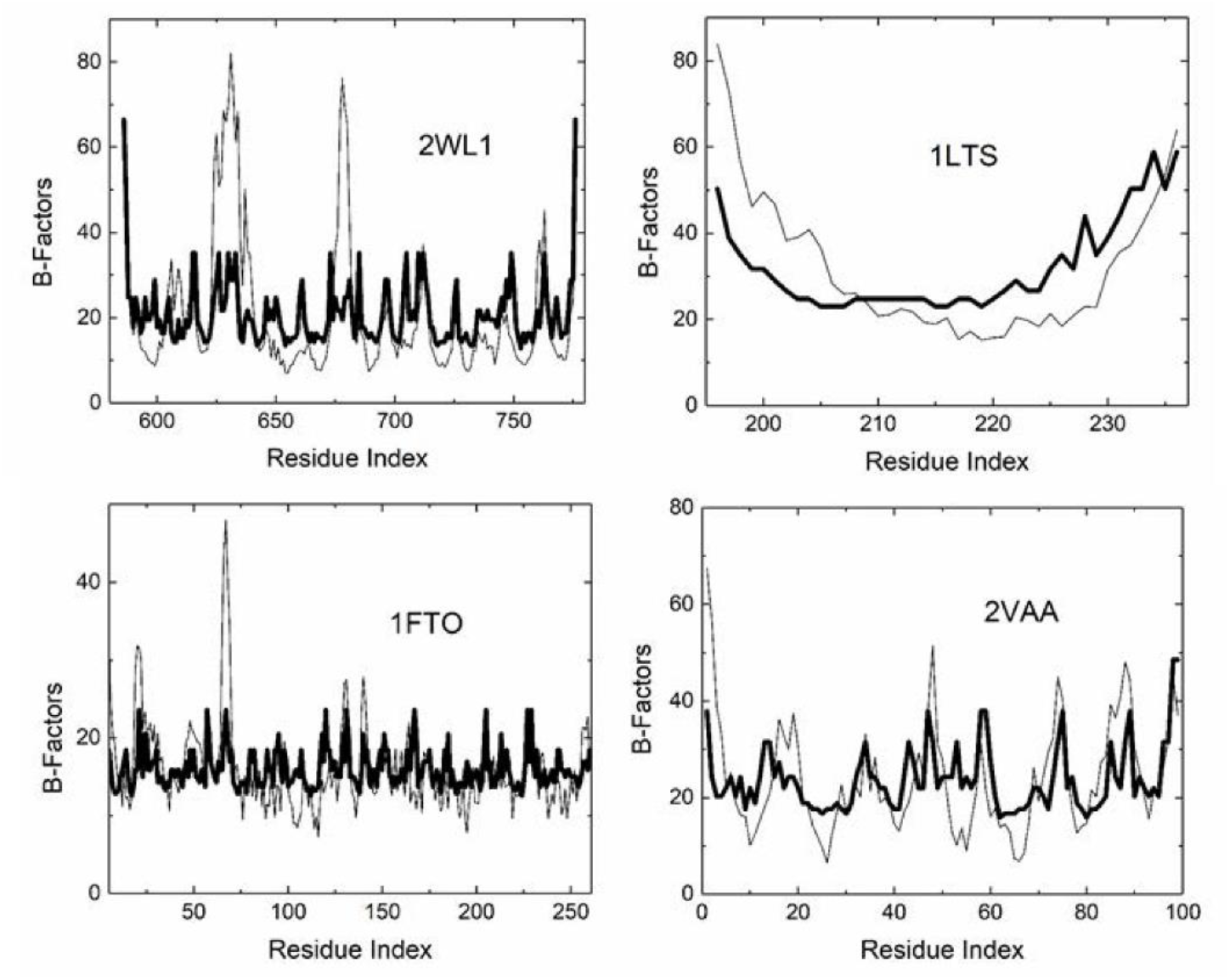
Comparison of experimental B-factors (thin lines) with the crowding effect (thick lines). The crowding effect is calculated by counting the alpha carbon neighbors within a 7 Ångstrom radius for each alpha carbon, taking the reciprocal of this count, and then fitting it to the experimental data using least squares. Network connectivity effects as obtained from the inversion of the Kirchoff matrix, for example, is not included in this comparison. The Protein Data Bank code of the proteins is shown in each panel.

Comparative analysis between GNM predictions and experimental B-factor profiles shows that despite its general efficacy, GNM systematically underpredicts loop or turn fluctuations while overpredicting fluctuations in densely packed regions. The underestimation of loop mobility can be attributed to several factors. First, the harmonic interaction assumption imposes a quadratic energy landscape, whereas loops often exhibit large-amplitude, anharmonic motions, particularly in flexible or disordered regions. Second, GNM’s fixed cutoff distance (typically 7–10 Å) for residue connectivity may inadequately capture the sparse contact patterns characteristic of extended loops. Third, long-range interactions become proportionally more significant in these sparsely connected regions, as they face less competition from the numerous short-range contacts that dominate in densely packed structural elements. Most significantly, loop dynamics are substantially influenced by entropic effects as these regions sample multiple conformational states—an aspect not explicitly addressed in the harmonic GNM framework.

Conversely, the over prediction of fluctuations in crowded regions suggests insufficient representation of multibody effects within GNM’s mathematical structure. A residue in the crowded environment of its neighbors may experience three body effects where a third residue may interfere with the interaction of i and j. Four body effects should also be considered where two residues k and l modify the neighborhood of i and j. These effects will be deeply affected by the size and type of side-chains and will modify the backbone coupling, electronic interactions and long-range correlations that emerge from the entire protein structure. The GNM has no context beyond counting number of neighbors and incorporating effects of chain connectivity on B-factors—a limitation originating from its theoretical roots in CJM of rubber elasticity, where interpenetration of junctions provides sufficient description for homogeneous polymeric systems, but becomes inadequate for proteins where residue diversity and specific interaction patterns demand contextual treatment. However, as noted above, B-factors reflect a composite of factors beyond neighbor and connectivity effects: secondary structure, solvent exposure, and sequence context, dynamic heterogeneity in the protein and multiple conformations and disorder. B-factors are the experimental outcome of these effects. A realistic model needs to be contextualized by systematically addressing these additional factors but it should contain the essential details avoiding unnecessary complexity yet be evolvable. An information-theoretic approach offers a promising avenue, employing concepts such as mutual information conditioned on the presence of several alpha carbons within the coordination shell of a given alpha carbon. This conceptual framework enables us to incorporate conditional dependencies and higher-order correlations such as three-body and higher correlations that may be crucial for accurately modeling the complex dynamics of protein structures.

The simplest refinement of the GNM is to keep the structure of the connectivity matrix (the Kirchoff matrix) unchanged while modifying the nature of the interactions and including those effects beyond network connectivity and interpenetration. This will modify the interpretation of the fictitious spring concept, which is rooted in the statistical mechanics of a Gaussian polymer chain, which fluctuates under forces applied to its ends[17]. The modification of the Kirchoff matrix should however be made in a self-consistent manner: The zero sum of each row should be retained to ensure equilibrium, the symmetry should be retained to preserve microscopic reversibility or detailed balance, and since every modification of the matrix also modifies its inverse covariance matrix, the iteration of this cycle should converge[18].

While our proposed method offers an approach to predict protein correlations with improved accuracy, the primary aim of this paper is to enhance the GNM by addressing factors not considered in its original formulation. We improve both the accuracy of B-factor predictions and the evaluation of covariances, the off-diagonal terms of the Kirchhoff inverse, in a self-consistent manner following the principles of information theory. Although accurate B-factor assessment is essential for understanding protein dynamics, proper evaluation of covariances is even more critical, as key functional mechanisms, such as allosteric signaling and energy transfer, propagate through residue correlations. Covariances are presently widely studied through time-intensive molecular dynamics simulations[19], with direct experimental determination only recently emerging[5, 6], making our computationally efficient approach particularly timely. The primary contributions of this work are threefold: we demonstrate significantly improved agreement between predicted and experimental B-factors compared to the standard GNM; we introduce a framework for contextualizing residue interactions within the GNM, enabling a more accurate representation of the complex correlation patterns that underlie protein function and allostery; and we develop a Monte Carlo approach to identify cross-correlations that satisfy the maximum entropy principle, providing new insights into how information is transferred within protein structures and how distant sites communicate to regulate function.

## 2. Materials and methods

### 2.1. The GNM and the formulation of the improved model

The ij’th element of the Kirchoff matrix of the original GNM is defined as

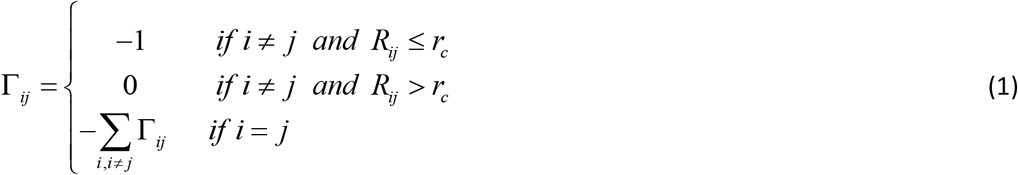

where *r*_*c*_ is the cutoff separation defining the range of interaction of non-bonded α carbons, and *R*_*ij*_ is the distance between the *i*th and *j*th *C*^*α*^ atoms. The presence of -1 at position (i, j) signifies the influence of residue j on the fluctuations of i. Since the interaction is inherently symmetric, the influence of i on j is identical, ensuring that the Kirchhoff matrix remains symmetric.

In the GNM, interactions lack context, with all contacts assumed to be identical and modeled solely as uniform pairwise interactions. However, pairwise interactions in proteins depend on multiple factors, including steric interactions, hydrogen bonds, and electrostatic forces, which introduce variability not captured by this simplification. Furthermore, the GNM overlooks higher-order interactions, such as triplet contributions—where the combined effect of neighbors j and k influences the fluctuations of residue i—or quadruplet interactions involving neighbors j, k, and l, which may also be modulated by these factors. Given that the interaction domain of a residue i may include up to 15 neighbors, these triplet and higher-order interactions could be significant. For instance, the presence of a pair k and l might expand the volume in which j fluctuates, thereby increasing j’s conformational freedom and reducing its impact on i. Such effects are not accounted for in the GNM, which thus serves as a zeroth-order approximation to the more refined model presented below.

The improved model retains the framework of the GNM while accounting for modifications to the i,j-potential based on the effects outlined in the preceding paragraph. Specifically, the interaction between residues i and j, beyond the uniform GNM potential, is adjusted to reflect the influence of other residues within the interaction domain of i and j. These modifications are incorporated into the i,j-th element of the Kirchhoff matrix, Γ_*ij*_, capturing the contextual and multiplet contributions to the pairwise dynamics as follows:

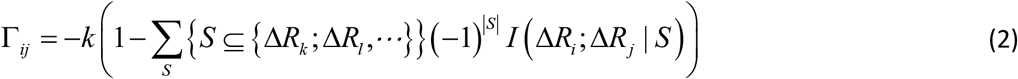

Here, k is the spring constant of the original GNM theory, the 1 in the parenthesis represents the spring force in the original GNM, and the sum represents the conditional mutual information between the fluctuations of residue i and j, systematically accounting for the effects of conditioning on the variables k, l, …. The summation runs over all subsets S of the set {Δ*R*_*k*_ ; Δ*R*_*l*_,}, meaning it considers all possible ways to condition on none, one, both, etc., of these variables. *I* (Δ*R*_*i*_ ; Δ*R*_*j*_ | *S*) denotes the mutual information between the fluctuations of residues i and j while conditioning on the subset S. The factor (−1) ^*S*^ alternates the sign depending on the number of elements in S. This ensures proper cancellation of redundant contributions, following the principle of inclusion-exclusion[20]. This correction represents a mean-field type adjustment, averaged over the fluctuations of all remaining residues k,l,… and will be explained in more detail below.

The summation term in Eq 2 represents the contributions of non-GNM effects to Γ. Inversion of this modified Γ leads to the covariance of fluctuations, which is then used to obtain the new correlation matrix Γ. We demonstrate that iterative application of this equation converges rapidly. At each step of the iteration, the Γ matrix evolves from the unweighted GNM representation to a dynamically reweighted interaction matrix that integrates information-theoretic corrections based on mutual information.

The diagonal elements of Γ^−1^ yield variances proportional to experimentally determined B-factors, providing a basis for validating the model’s accuracy. More importantly, the off-diagonal elements of Γ^−1^ show correlations between residue fluctuations. Since the inverse operator generates correlations between spatially distant residues, an accurately constructed Γ is crucial for understanding allostery and information/entropy transfer in proteins.

### 2.2. Detailed derivation of the improved Γ matrix

The summand in Eq. 2 expands to:

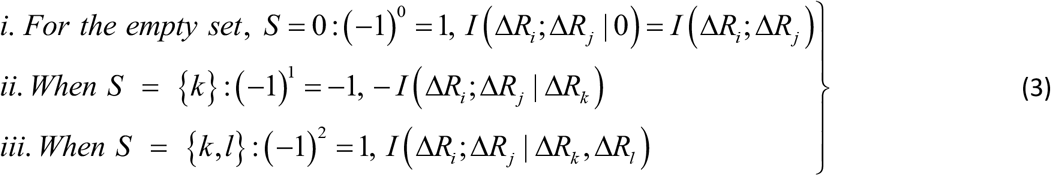

Where, *I* (Δ*R*_*i*_ ; Δ*R*_*j*_), *I* (Δ*R*_*i*_ ; Δ*R*_*j*_ | Δ*R*_*k*_), and *I* (Δ*R*_*i*_ ; Δ*R*_*j*_ | Δ*R*_*k*_, Δ*R*_*l*_) are the joint mutual information between residues i and j, the mutual information between i and j after accounting for the influence of residue k, and the mutual information after accounting for the joint effect of residues k and l, respectively. We consider only up to four body interactions. Using Eqs. 3 in Eq. 2 and properly scaling each term we obtain

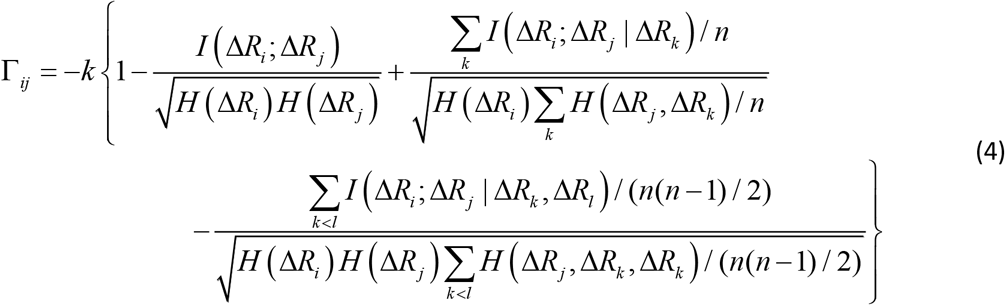

where, *H* (Δ*R*_*i*_), *H* (Δ*R*_*j*_, Δ*R*_*k*_) and *H* (Δ*R*_*j*_, Δ*R*_*k*_, Δ*R*_*l*_) are the singlet, doublet an triplet entropies and n is the number of neighbors of i. This systematically accounts for the information between residues i and j while properly including and excluding the effects of residues k and l. The alternating signs ensure that there is no double-counting information and are correctly accounting for how k and l individually and jointly affect the i-j interaction. In Eq 4, in order to align with GNM’s unit contact strength -1, Γ_*ij*_ is refined using averaged entropy terms, 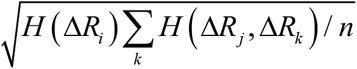 and 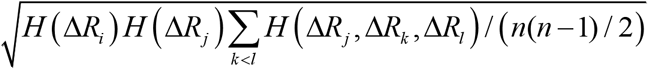. These factors ensure adjustments that are commensurate with the GNM’s unit contact strength (−1), typically yielding values below 1, which preserves matrix stability and avoids over-amplification. An alternative normalization to [0, 1] using 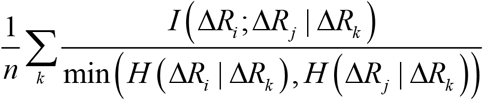 and 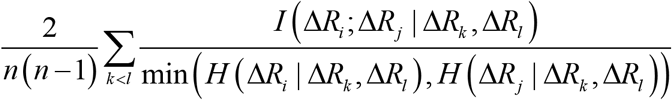 was tested but introduced negative eigenvalues, confirming the original scaling’s suitability.

Eq. 4 will be evaluated iteratively where the zeroth iteration will be the GNM. The mutual information terms are functions of the covariance of the fluctuations. The covariance obtained by inverting Γ will be used to obtain the new mutual information values, which will be used on the right hand side in the next iteration. Iterations of Eq 4 give the modified GNM. The doublet correction, ij term, in Eq 4 decreases the constraining effect of j on i. This decrease has an increasing effect on the fluctuations of i. The triplet correction, ijk term, increases the constraint on i. This is the leading term of the crowding effect, which suppresses the fluctuations of i in a crowded environment. The quadruplet correction, ijkl term, decreases the effect, which is a dilution effect due to the presence of two residues around i.

### 2.3. Aim and Strategy of the Improved GNM

The improved GNM aims to enhance protein dynamics predictions by optimally partitioning residues into three sets, Set 0 *I* (Δ*R*_*i*_ ; Δ*R*_*j*_), Set 1 *I* (Δ*R*_*i*_ ; Δ*R*_*j*_ Δ*R*_*k*_), and Set 2 *I* (Δ*R*_*i*_ ; Δ*R*_*j*_ Δ*R*_*k*_, Δ*R*_*l*_), to maximize agreement with experimental B-factor profiles. Unlike the standard GNM’s uniform interactions, our approach assigns residues to sets based on their dynamic roles, with Set 1 and Set 2 sharing identical residues to model three- and four-body interactions with alternating signs, per the inclusion-exclusion principle. The core challenge is to allocate large B-factor residues, typically in flexible loops or turns to Set 0 for pairwise interactions, and small B-factor residues, often in crowded cores, to Sets 1 and 2 for multi-body corrections, optimizing the Kirchhoff matrix (Eq. 4) for accurate fluctuations. While most assignments align with this dynamic profile, our analysis shows exceptions where residues are partitioned into unexpected sets, e.g., residues in sparse environments into Set 1/Set 2 or residues in crowded neighborhoods into Set 0. These anomalies illuminate the complexity of protein dynamics beyond local density, as evidenced by outliers in our neighbor count versus inverse B-factor scatter (Figure 1).

Several factors explain these exceptions. Residues in secondary structures, such as helix termini, may exhibit low B-factors but be assigned to Set 0 if they drive flexible pairwise interactions with adjacent loops. Conversely, surface residues with high B-factors may join Set 1 when solvent-mediated or multi-body constraints limit their fluctuations, reflecting effective rigidity. Allosteric residues, critical to long-range cooperativity, may assemble into networks that favor information flow rather than following conventional B-factor distributions. Additionally and most importantly, allosteric effects within the coordination shell may assign residues to sets that balance system-wide dynamics, favoring collective agreement over local B-factor fidelity. These exceptions indicate the need to contextualize assignments, linking them to structural features like secondary structure, solvent exposure, or allosteric roles, thus improving our understanding of protein function and dynamics.

In the analysis below, we occasionally use the metric percent solvent accessibility, %SAS, of a residue which is calculated by first determining the absolute solvent-accessible surface area (ASA) for that residue, using the Lee and Richards method, which models a water molecule as a sphere (radius 1.4 Å) rolling over the protein surface[21]. The ASA is then compared to a reference maximum value for that residue type—usually the surface area of the same amino acid in an extended tripeptide conformation (Gly-X-Gly, where X is the residue of interest). Due to this definition, percent solvent accessibility may exceed 100%. It often explains the unexpected deviation of B-factors from an expected value in contextualizing the model.

To achieve the optimal partitioning, we employ a Monte Carlo approach: residues are ordered by decreasing B-factors, iteratively added to Set 0 (large B-factors) or Sets 1 and 2 (small B-factors), and retained if the RMSD decreases or removed if it increases, until convergence. Post-convergence, we apply the maximum entropy (MaxEnt) principle to derive covariance matrices that maximize uncertainty with four constraints: (i) the optimized B-factors are kept fixed, (ii) connection topology is fixed, i.e., zero elements of the Kirchoff matrix remain zero, (iii) symmetry of the matrix is retained, (iv) zero sum row property is retained. This yields robust predictions without adjustable parameters. We then contextualize each residue’s set assignment, explaining its role based on local density, secondary structure, solvent exposure, or allosteric connectivity, as evidenced by enhanced entropy transfer in allosteric pathways. This partitioning and contextualization framework offers an interpretable model for protein dynamics. Importantly, the proposed method is not intended to predict B-factors or covariance matrices but to clarify the contributions of factors, such as local density, secondary structure, solvent exposure, and allosteric connectivity, not accounted for in the original Gaussian Network Model (GNM), thereby enhancing our understanding of the underlying physics.

### 2.4. Evaluation of the cross-correlations from B-factors, Maximum entropy formulation

For a protein with n residues, matching the diagonal elements of Γ^−1^ to B-factors constrains only n values, leaving the n(n−1)/2 off-diagonal elements of Γ, representing inter-residue covariances, underdetermined. To resolve this, we apply the maximum entropy principle, maximizing the system’s configurational entropy *S* = log (det(Γ^−1^)) (excluding the zero eigenvalue for translational invariance) while constraining the optimized B-factors. Starting from a mutual information-refined Γ, incorporating mutual information terms (e.g., *I* (Δ*R*_*i*_ ; Δ*R*_*j*_ Δ*R*_*k*_)), we use Monte Carlo optimization to adjust off-diagonal elements. The MC scheme enforces: (1) symmetry Γ_*ij*_ = Γ _*ji*_, (2) zero row sums 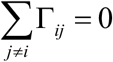, and (3) alignment of Γ^−1^ diagonals with B-factors via least-squares fitting. At each step, we perturb a randomly selected off-diagonal Γ_*ij*_ (within the initial topology) by δ ∼ Uniform(−0.1,0.1), Γ_*ij*_, Γ _*ji*_, Γ_*ii*_

We compute Γ^−1^, calculate the RMSD between its diagonal and B-factors, and evaluate the cost function *C* = log(det (Γ^−1^))+ *λRMSD*^2^, with λ=10^4^ balancing entropy and accuracy.

A Metropolis criterion accepts perturbations that reduce C or with probability exp(−Δ*C* / *T*), where T cools linearly from 1.0 to 0.01 over 100,000 steps. Optimization stops when the given maximum number of iterations is reached. This yields a physically meaningful covariance matrix, enhancing the GNM’s ability to predict dynamic correlations.

### 2.5. Mutual information and entropy transfer in terms of Gaussian variables

Several measures of allosteric communication have been proposed[22-28], among these, mutual information and entropy transfer are the most significant measures. In this section, we expand the formal expressions presented above in terms of Gaussian variables and discuss mutual information and entropy transfer in terms of these.

For a multivariate Gaussian system, mutual information is given in terms of mean-squared fluctuations of residues by[23]

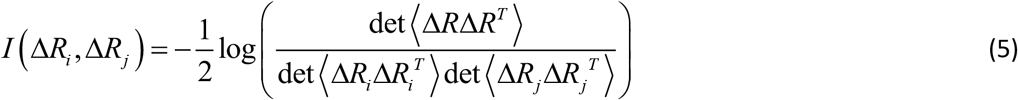

where Δ*R* is the 6×1 column vector Δ*R* = *col* (Δ*X*_*i*_, Δ*Y*_*i*_, Δ*Z*_*i*_, Δ*X* _*j*_, Δ*Y*_*j*_, Δ*Z* _*j*_,) and the superscript T denotes transpose. For GNM which assumes isotropic fluctuations, Eq. 5 takes the following form:

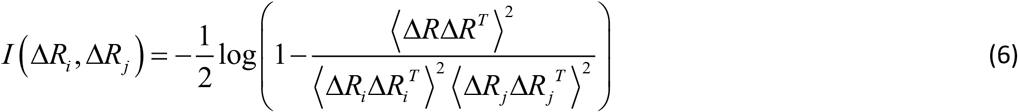

Mutual information given by Eq. 5 measures how much knowing the fluctuation of residue i tells us about the fluctuations of residue j. Summing over j gives the cumulative mutual information, CMI,

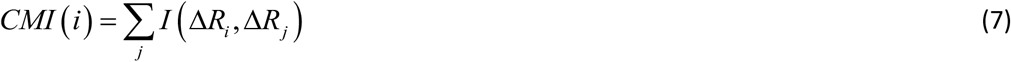

CMI quantifies how much the fluctuations of residue *i* are statistically linked to the fluctuations of the rest of the protein. A high CMI means that residue *i* shares a lot of mutual information with many other residues—it is informationally influential in the global dynamics. CMI captures the total dynamic coupling of residue *i* with the rest of the protein, making it a natural candidate for identifying residues involved in allosteric regulation or transmission.

Entropy transfer *T*_*i*→ *j*_ is defined as a measure of the reduction in uncertainty (or equivalently, the gain in information) about the future state of residue j given the past state of residue i, conditioned on the past state of j. We will be using entropy transfer and information transfer interchangeably. Thus, information transfer from residue i to residue j with a lag time *τ* is defined as[29-32]

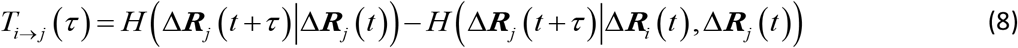

where, the first and second terms on the right hand side are the conditional entropies. For a multivariate Gaussian system, with Δ***R*** denoting the column vector of residue fluctuations and Δ***R***^*T*^ showing its transpose, entropy is written as

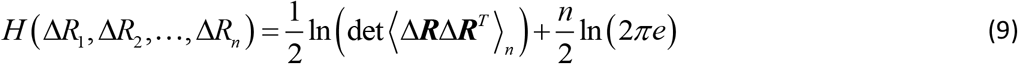

where in this case Δ*R* = *col* (Δ*X*_1_, Δ*Y*_1_, Δ*Z*_1_,, Δ*X*_*n*_, Δ*Y*_*n*_, Δ*Z*_*n*_,). Conditional entropies may be derived from Eq. 9 using known relations[28]. Substituting into Eq. 5 leads to the explicit form[31]

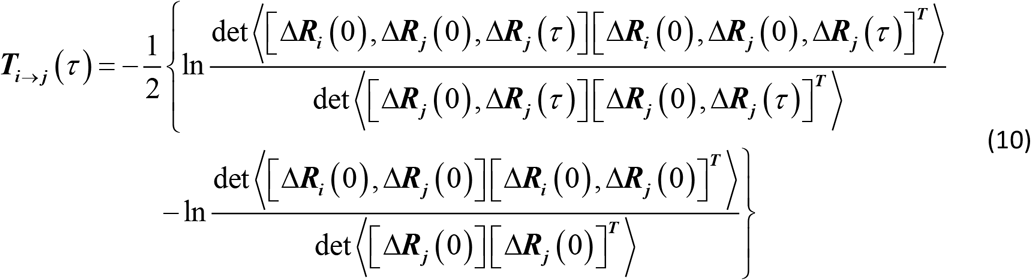

When the determinants are expanded, there are terms of the form ⟨Δ***R***_i_(0),***R***_j_ (τ) ⟩ which are evaluated[33] according to the expression

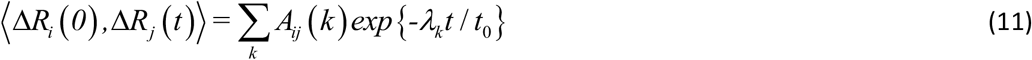

where,

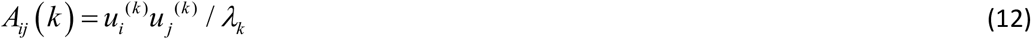

with *λ*_*k*_ being the kth eigenvalue and 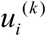being the ith component of the kth eigenvector of the Kirchoff matrix.

The standard GNM captures protein dynamics through entropy transfer, the propagation of information between residues, which mediates functional motions such as allosteric signaling[31, 34]. Our improved GNM, incorporating mutual information corrections to the Kirchhoff matrix (Eq. 4), refines these dynamics by accounting for multi-body interactions and B-factor-derived classifications. To assess the impact, we recalculated mutual information and entropy transfer for allosterically known proteins.

## 3. Results

### 3.1. Testing the hypothesis

The improved GNM rests on the hypothesis that residues in sparse environments, such as loops, turns, or coils, exhibit enhanced fluctuations compared to standard GNM predictions, whereas those in densely packed regions, like protein cores, show reduced fluctuations. However, local density alone does not fully account for these deviations, as shown by scattered outliers in our neighbor count versus inverse B-factor plot (Figure 1). Experimental B-factors, which correlate inversely with neighbor counts while capturing secondary structure, solvent interactions, nonlinear dynamics, and other influences, provide a comprehensive measure of residue fluctuations. Thus, we order residues by increasing B-factors, allocating high B-factor residues to Set 0 for pairwise interactions and low B-factor residues to Sets 1 and 2 for multi-body corrections, enhancing dynamic predictions.

In Table 1, we compare the RMSD between experimental and GNM predicted B-factors and the RMSD between experimental and improved GNM predicted B-factors. We tested several examples, and present results from eight representative samples in Table 1. The first column shows the PDB code of the protein. The second column is the RMSD between the experimental data and GNM B-factors. The third column is

**Table 1.** Deviation (RMSD) of the original and the improved GNM from experimental B-factors.

|  |  |  |  |
| --- | --- | --- | --- |
| Protein | RMSD<br>GNM | RMSD<br>improved GNM | percent improvement<br>over GNM |
| 6GOD | 8.33 | 5.09 | 38.9 |
| 1F4V | 5.84 | 3.61 | 38.2 |
| 1ABA | 3.17 | 1.80 | 43.2 |
| 1FTO | 4.54 | 3.06 | 32.6 |
| 2OMF | 12.65 | 7.24 | 42.8 |
| 2VAA | 8.82 | 4.74 | 46.3 |
| 2WL1 | 14.05 | 10.46 | 25.6 |
| 3K0N | 4.27 | 3.12 | 26.9 |

the RMSD between the experimental data and the improved GNM B-factors. Percent improvement, calculated as 100x(RMSDGNM-RMSDimprovedGNM)/RMSDGNM is presented in the fourth column. Calculations are made according to Eq. 4 where the pair interactions (second term in the braces in Eq.4) are calculated for Set 0 and the triplet and quadruplet interactions (third and fourth terms in braces in Eq. 4 are calculated for Set 1 and 2 residues. The model’s success, without adjustable parameters, hinges on ordering residues by B-factors to define these sets.

The calculations in Table1 are made for a cutoff length of 7.8 Å. However, the results are not strongly dependent on the choice of cutoff radii. B-factor convergence took place in 5 to 20 iterations, depending on the size of the protein. The residues in Sets 0, 1 and 2 are given in the Supplementary material, section S-1.

### 3.2. Changes in B-factors, correlations, mutual information and entropy transfer: Analysis of wild type KRAS

In this section, we evaluate the changes in various variables resulting from B-factor optimization and entropy maximization, using the widely studied protein KRAS as an example. KRAS is a 172 residue GTPase involved in regulating multiple signaling pathways through its interaction with various effector proteins, such as RAF, PI3K, and RalGDS. The two functional loops of KRAS are known as switch-I and switch-II. These regions undergo conformational changes upon GTP binding, enabling KRAS to interact with its effector proteins and regulators. Specifically, switch-I typically encompasses residues 30–40, while switch-II spans residues approximately 58–72. Together, these loops form the binding interface critical for KRAS activation and downstream signaling. The original GNM gives an RMSD of 8.33 with experimental B-factor values and the improved GNM gives an RMSD of 5.09, an improvement of 38.9% as may be seen from Table 1.

In Figure 3 we compare the experimental B-factor profile of 6GOD.pdb (thin line) with the results of the improved GNM, thick line. The improvement predicts all major peaks and dips and predicts the details with some accuracy. The contextualization residues are indicated with hollow and filled circles and discussed below.

**Figure 3.**
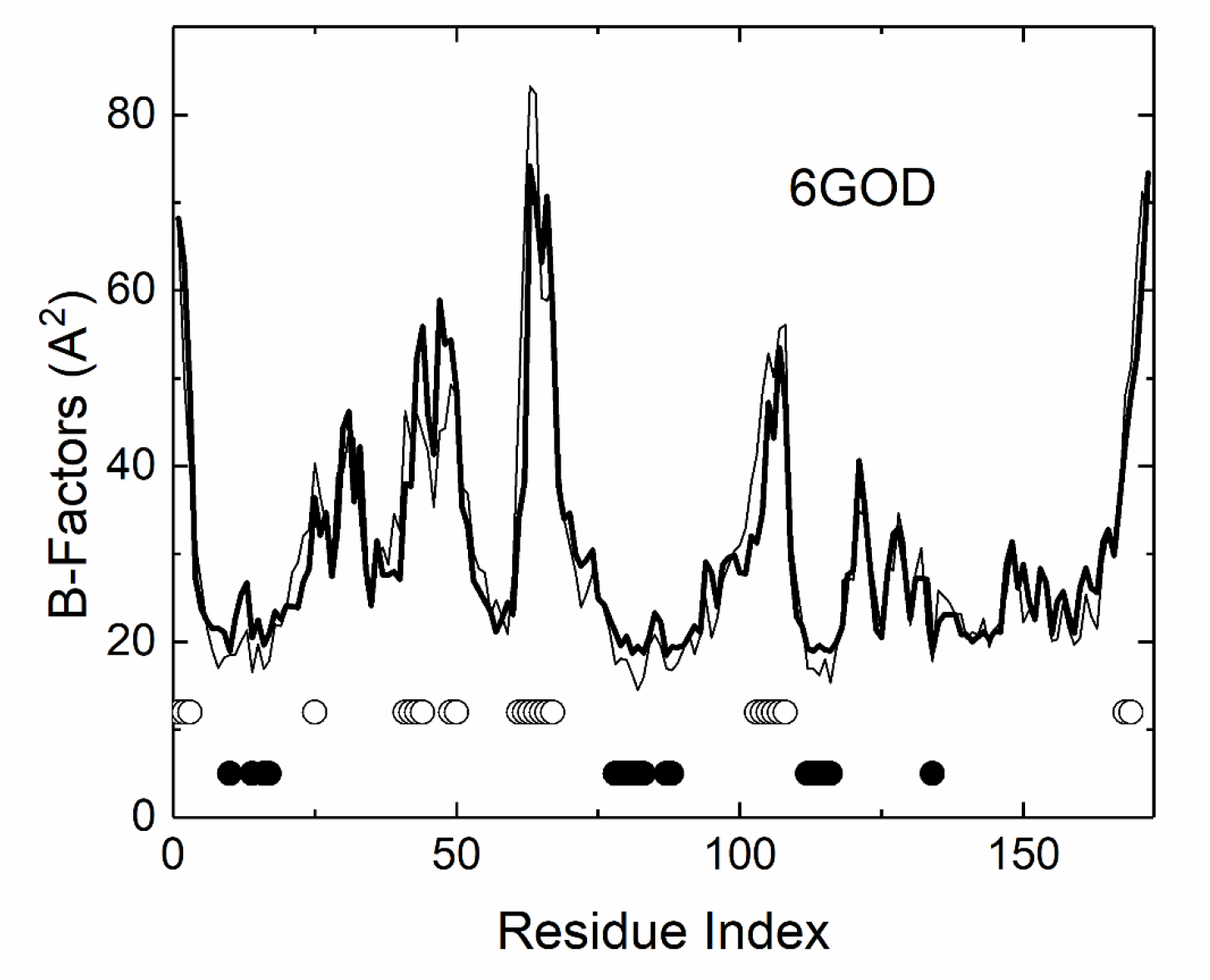
Comparison of experimental (thin line) and improved GNM predicted B-factors. The residues of Set 0 are shown with unfilled circles and those of Set 1 and 2 are shown with filled circles.

Residues whose effects are not sufficiently accounted for the original GNM are shown by the circles. Residues for pair interactions, Set 0, are shown with hollow circles. Residues with triple and quadruple interaction effects, Sets 1 and 2, are indicated with filled circles.

**Set 0** consists of residues with high mobility not sufficiently accounted for by the original GNM. This includes N-terminal residues 1–3 and residue 25, which lies at the end of a helix and displays 54% solvent accessibility (see *Materials and Methods* section for definition). Notably, residues 41–44, which belong to a β-strand, also show unexpectedly high mobility. This may be due to their elevated solvent accessibility—66% for Arg41 and 79% for Gln43. Interestingly, Switch I (residues 30–40), a key component of the effector-binding interface, exhibits lower mobility than its flanking hinge regions (See further explanationbelow). Additional high-mobility residues include 49 and 50, located at a β-turn–β motif; residues 61–67, which make up the Switch II loop and include the allosterically important Gln61; the helix–turn–β-strand motif at residues 103–108; and residues 168–169 at the C-terminal helix end, located near the hypervariable region responsible for membrane localization.

**Sets 1 and 2** include residues with lower mobility than that predicted by the original GNM, many of which are structurally buried and likely play stabilizing roles. For example, residue 10, with 0% solvent accessibility, serves as a stationary hinge point connected to a loop containing the critical residues 12 and 13. Residues 14, 16, and 17, also with 0% solvent accessibility, lie within a β-loop–helix motif.

Residues 78–83, likewise with 0% solvent accessibility, are situated on a β-strand flanked by two other β-strands, contributing mechanical rigidity and structural integrity to the overall structure of KRAS. Residues 87–88 lie on a β-turn–helix motif and show high solvent accessibility (51% and 110%, respectively); however, their low B-factor values are attributed to high local contact densities (neighbor counts of 12). Residues 112–116 form part of a central β-strand within a β-sheet and are fully buried (0% SAS). Finally, residue 134 is centrally located within a helix that faces the β-sheet.

Entropy maximization is performed on the B-factor improved results according to the principles explained in the theory section. 100,000 MC steps are sufficient for convergence as shown in Figure 4. The code used for calculating improved B-factors and MaxEnt is available on request. The parameters used for calculations are given in the code. All quantities used in the figures are based on the dimensionless form of the Shannon equation and its derivatives. Multiplying the Shannon equation for entropy and the information theory expressions derived from it with kBT, where kB is the Boltzmann constant and T is the absolute temperature, gives the free energy equivalent of entropy which we use below to compare results with known thermodynamic quantities.

**Figure 4.**
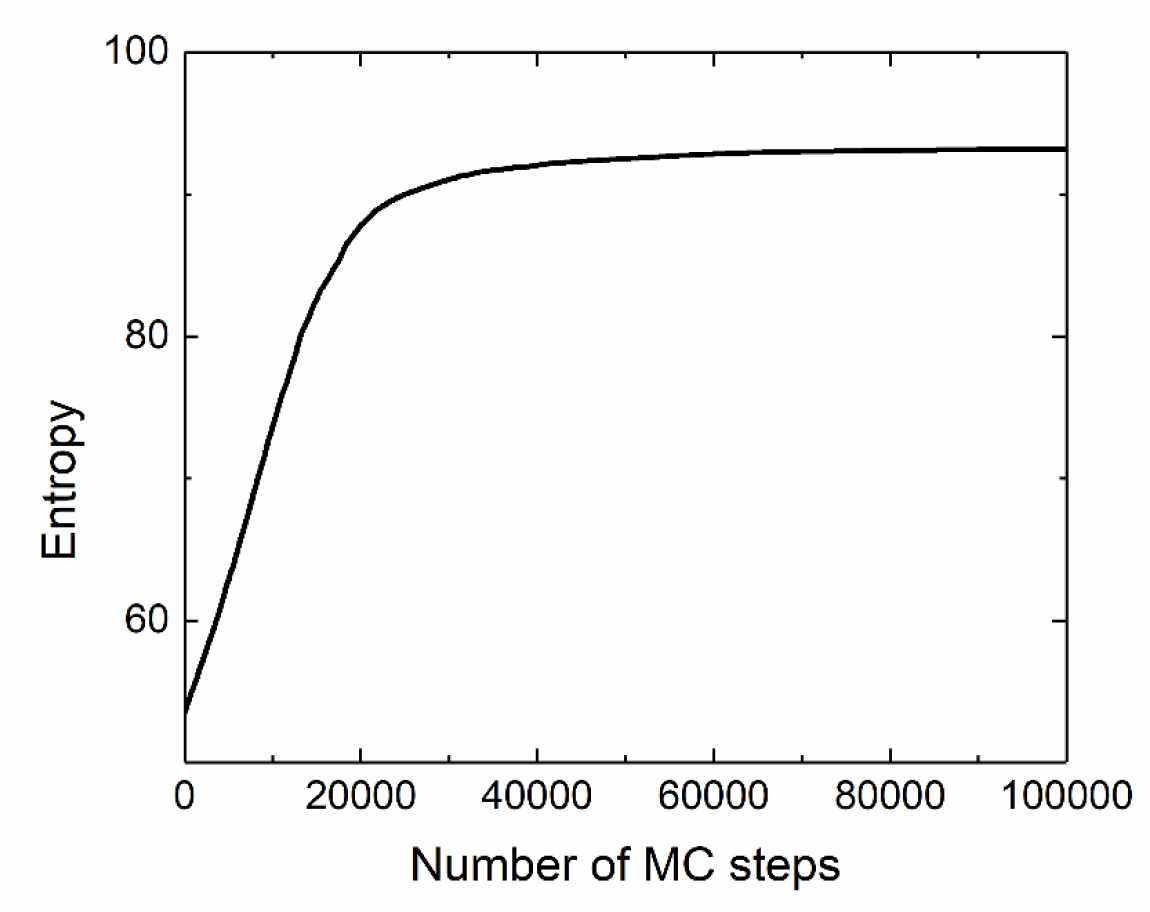
Increase and convergence of entropy as a function of MC steps.

A wide range of parameters is tried and the ones leading to stable and reproducible results are used. Using Eq. 9, and kBT=2.49 kJ/mol, the starting entropy of the Kirchoff is 132 kJ/mol and the maximum converged entropy is 234 kJ/mol. On per residue basis, these correspond to 0.77 and 1.36 kJ/mol, which appear to be in the right ballpark with known thermodynamic parameters for protein fluctuations, noting that our model captures only translational motions of the alpha carbons.

Results for cumulative mutual information, CMI, for the GNM and entropy maximized GNM are shown in Figure 5. Comparison shows that the shapes of the profiles do not differ from each other significantly. Entropy maximization increases the CMI values between residues 60-80 and 110-150.

**Figure 5.**
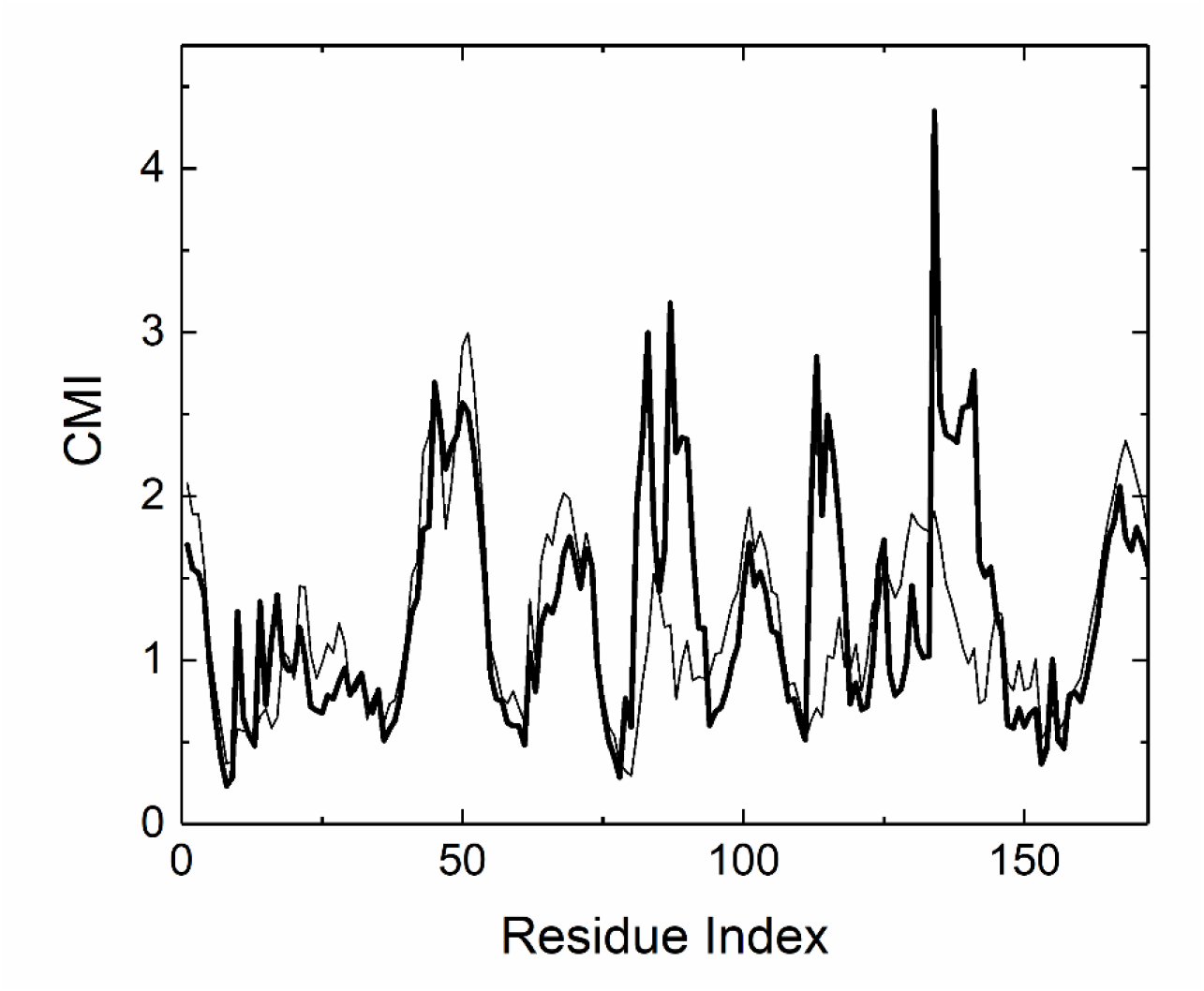
CMI profile for entropy maximized GNM (thick line) and original GNM (thin line).

In the GNP-bound KRAS structure (6GOD.pdb), Switch 1 and Switch 2 residues, such as Y32 and Q61, exhibit low B-factors, indicating restricted mobility, and are located in the minima of the cumulative mutual information (CMI) profile, reflecting weak dynamical correlations with other residues. This behavior arises from three factors: (i) Strong local constraints with low global embedding: These residues are stabilized by interactions with GNP, magnesium, and potentially effector or inhibitor molecules, resulting in low B-factors and limited motional freedom. Their motions are weakly correlated with the rest of the protein, leading to low CMI, as they act as functional toggles responding sharply to local signals (e.g., GTP hydrolysis or effector binding) rather than participating in globally coordinated motions. (ii) Functional independence: Their dynamical decoupling enables conformational changes critical for GTP hydrolysis (Q61) and effector binding (Y32), allowing independent responses to functional cues. (iii) Entropic decoupling: These functionally active regions are entropically decoupled from the global correlation network, facilitating conformational switching essential for KRAS’s allosteric activity.

Mutual information for residues 87 and 134 are compared for GNM and improved GNM in Figure 6. The abscissae are the other residues. The improved GNM gives larger amplitudes compared to those of the GNM values.

**Figure 6.**
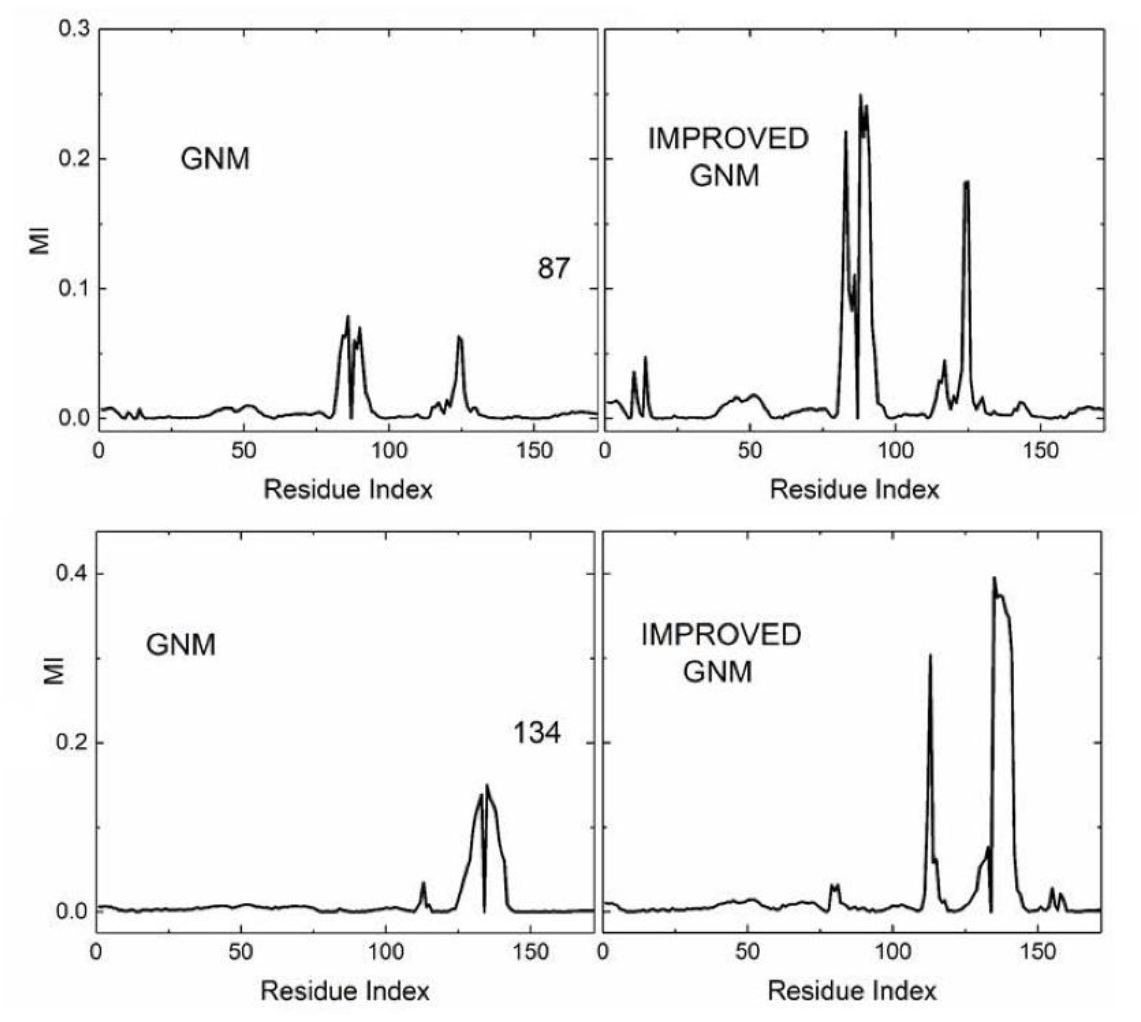
Mutual information for residues 87 and 134 using the original GNM (top and bottom left panels) and the improved GNM (top and bottom right panels)as a function of all other residues of the protein.

Entropy transfer between pairs of residues to which the guanine nucleotides bind are shown in Figure 7. Unlike the B-factors, correlations and mutual information, where Kirchoff matrix of GNM and entropy maximized GNM give results of the same order of magnitude, entropy transfer values obtained from the entropy maximized Kirchoff may be significantly different than those of original GNM. Static properties (B-factors, correlations) primarily capture equilibrium behavior whereas entropy transfer depends on the specific pathways and timing by which information flows. Modifying the Kirchhoff matrix likely disrupts these specific pathways disproportionately. Also, entropy transfer typically involves derivatives or differences of conditional entropies and these higher-order calculations tend to amplify small changes in the underlying model where a moderate reduction in coupling strength can cause a non-linear (exponential) decrease in entropy transfer. Finally, the modified GNM preserved the overall connectivity pattern but involves reduced connection strengths. Comparison of the upper and lower panels of Figure 7 shows that the improved GNM may lead to up to two orders of magnitude smaller transfer values. The right upper and lower panels show that the directions of transfer may also be different.

**Figure 7.**
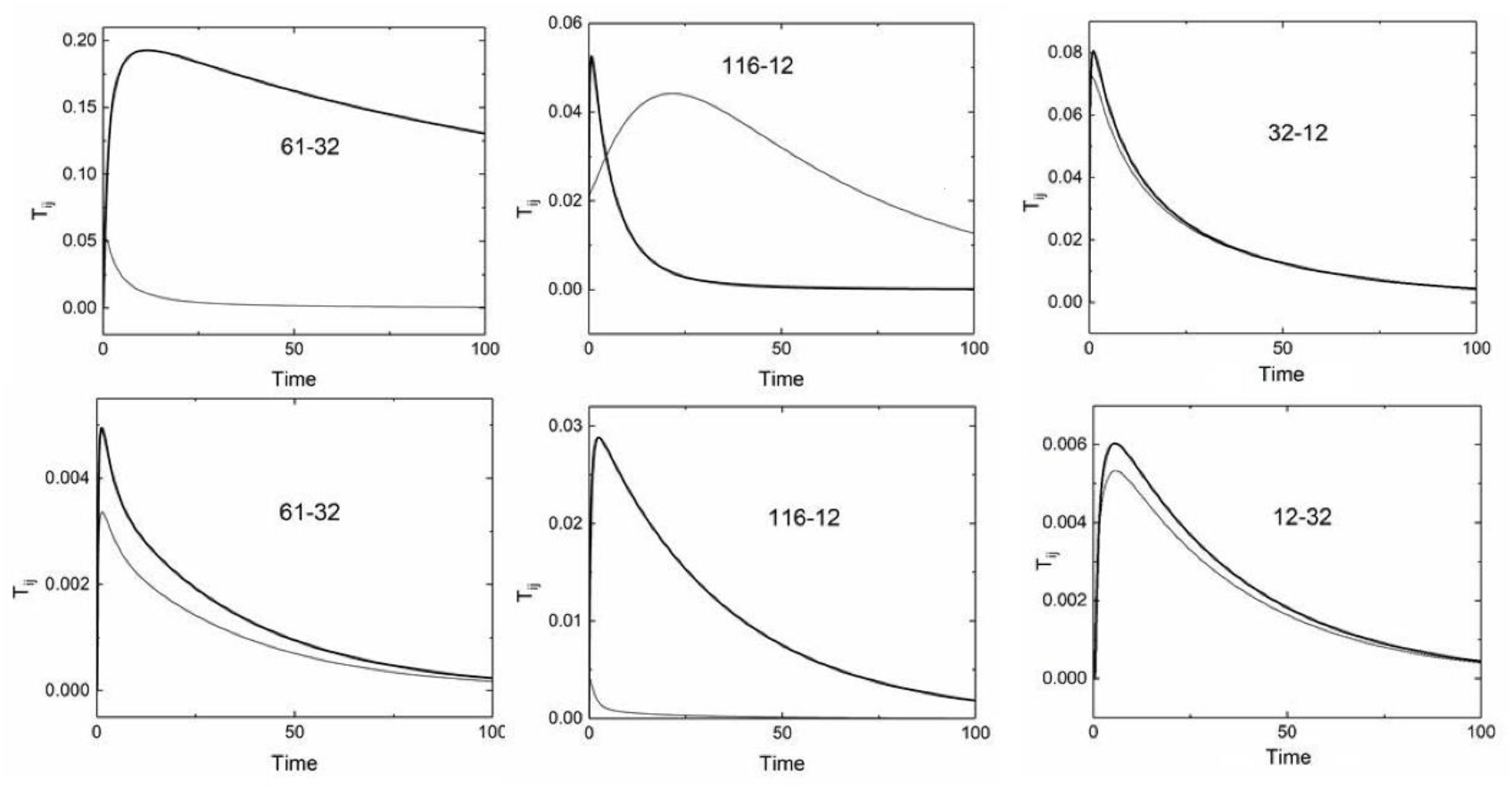
Entropy transfer decay curves for various pairs of residues. The top three panels are obtained using the original GNM. The lower three panels are obtained from improved GNM after MaxEnt. Thick lines show the dominant direction identified on each curve. Light curve is transfer in the opposite direction. Calculations are performed according to Eq. 4.

The dominant direction in Figure 7 is shown with the thick lines. The units of entropy and time are not indicated in the figure because the equations are treated as dimensionless quantities in the model[31, 33].

KRAS has several drug binding pockets[35] which play roles in the interaction of the drug with the residues of the protein. Noteworthy among these are the residues 17, 32, 95, 99, 107. In Figure 8 we show entropy transfer plots that show the role of these residues as either information donor or receptor points in the protein. The heavy lines in the figure indicate the direction of entropy transfer. Accordingly, information is transferred from the nucleotide binding residues 11 and 12 to 32 and 95, respectively; from the hydrophobic pocket residue 72 to the α2–α3 interface residue 99 and then from 99 to the P110 site residue 107; from allosteric site 3 residue 17 to the Switch 1 residue 28; from the α2–α3 interface residue 95 to the cysteine residue 51. The 10-fold increased decay time from 95 to 51 in the lower right panel of Figure 7 is worth noting. Strongest information transfer is between two contacting residues, 72 and 99, of Helices 2 and 3.

**Figure 8.**
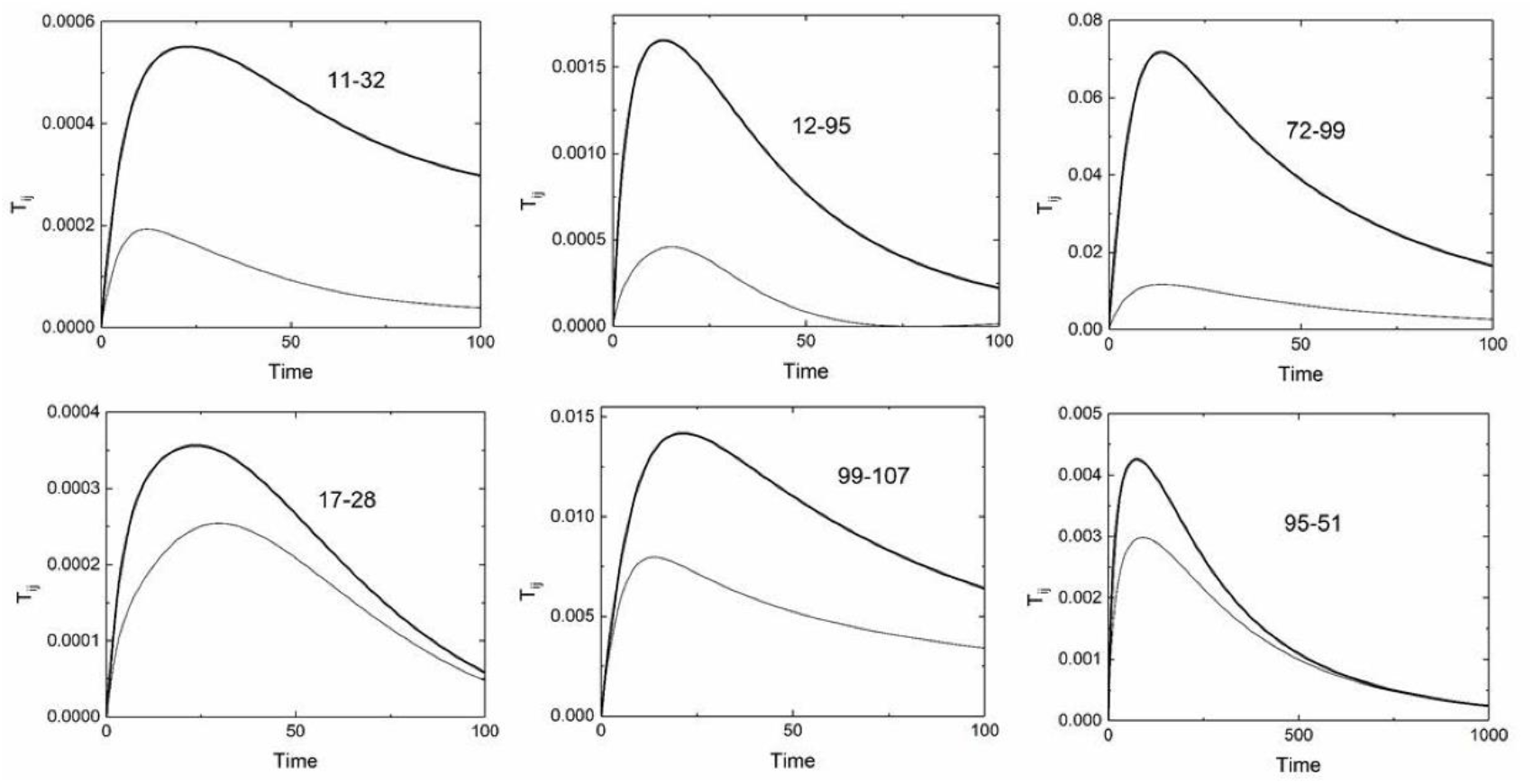
Directional dependence of information flow between allosterically known residues. Labels of residue pairs in each panel correspond to the thick lines. Thin lines are for flow in the reverse direction.

While direct experimental data on the direction of information flow in KRAS are lacking, the entropy-maximized model offers a framework for predicting these directions. Its reliability derives from several key attributes. By constraining the model to reproduce experimentally derived B-factors while maximizing entropy, it incorporates empirical data while minimizing artificial correlations unsupported by experiment. This approach adheres to the principle of maximum entropy, ensuring minimally biased inferences based on available constraints. Additionally, the model accounts for many-body interactions through conditional probabilities, more accurately capturing the cooperative nature of protein dynamics compared to models relying solely on pairwise correlations. By integrating statistical mechanical principles with experimental observations, the entropy-maximized model generates robust predictions of directional information flow, providing a foundation for future experimental validation.

## 4. Discussion

The GNM has been successful in advancing our understanding of protein dynamics, providing a computationally efficient framework to model proteins as elastic networks of alpha carbons connected by harmonic springs. However, its reliance on uniform pairwise interactions and a fixed cutoff distance limits its accuracy in capturing the complex interplay of forces governing residue fluctuations. The improved GNM presented here addresses these limitations by incorporating information-theoretic corrections to the Kirchhoff matrix, accounting for contextual and higher-order interactions. This results in improved B-factor predictions, with RMSD reductions of 25.6–46.3% across eight representative proteins (Table 1), and more accurate covariance matrices that better reflect the cross-correlations critical for allostery and information transfer.

The standard GNM systematically underpredicts fluctuations in flexible loops and overpredicts them in densely packed regions due to its harmonic assumptions and lack of multi-body interaction terms. Our approach helps improve these issues through a dynamically reweighted Kirchhoff matrix (Eq. 4), adjusted using conditional mutual information to capture triplet and quadruplet effects. The iterative optimization converges rapidly (5–20 iterations), balancing local and long-range contributions, while the Monte Carlo-driven maximum entropy approach refines the covariance matrix by maximizing configurational entropy under experimental B-factor constraints. This ensures the least biased estimate of residue correlations, minimizing assumptions unsupported by data.

A key strength of the model is its contextualization of residue dynamics. By partitioning residues into Set 0 (high B-factor, pairwise interactions) and Sets 1 and 2 (low B-factor, multi-body interactions), it captures nuanced roles based on structural and dynamic environments. In KRAS (PDB: 6GOD), flexible regions like Switch I and Switch II are assigned to Set 0, reflecting their high mobility, while buried β-sheet or helical residues are assigned to Sets 1 and 2, accounting for crowding effects. Exceptions, such as solvent-exposed residues in Sets 1 and 2 or buried residues in Set 0, show the limitations of local density metrics (Figure 1) and highlight the influence of secondary structure, solvent accessibility (%SAS), or allosteric connectivity. These findings point to the need for a systematic approach to residue assignment, integrating multiple structural features.

The model’s predictions of mutual information and entropy transfer provide insights into allosteric communication. In KRAS, increased cumulative mutual information (CMI) for residues 80-93, 110–118, and 133-143 (Figure 5) highlights their role in effector binding and signaling, while low CMI in Switch I and Switch II suggests functional independence, enabling rapid conformational switching. Entropy transfer analysis (Figures 7 and 8) identifies directional information flow, such as from residue 95 to 51, with a 10-fold increased decay time in the improved model, aligning with known allosteric pathways.

However, these predictions rely heavily on B-factor agreement, as direct experimental data on cross-correlations or entropy transfer (e.g., from NMR or single-molecule spectroscopy) are scarce. While entropy maximization ensures a theoretically robust covariance matrix by minimizing biased complexities, the lack of additional experimental validation limits the model’s empirical grounding. Future studies should prioritize experimental techniques to verify predicted correlations and information flow, enhancing the model’s credibility.

## 5. Conclusion

This study presents an improved GNM that integrates information-theoretic corrections and maximum entropy principles, achieving better B-factor predictions and robust estimates of residue correlations, mutual information, and entropy transfer. By contextualizing residue interactions, the model overcomes the standard GNM’s uniform interaction assumption, offering a detailed view of protein dynamics with applications in understanding allostery and designing therapeutics. The application to KRAS illustrates its capability to study functional pathways, reinforcing its potential in computational biophysics.

A critical consideration is the indeterminacy in optimizing the Kirchhoff matrix’s off-diagonal elements, as the n(n-1)/2 covariance terms are underdetermined by n B-factors. Prior work by Erman[18] used a Monte Carlo approach to iteratively adjust spring constants for near-perfect B-factor agreement, achieving excellent correlations but drawing criticism for potential overfitting and lack of physical basis[36]. The concern, which is valid, is that empirical fitting may yield multiple Kirchhoff matrices with identical B-factors but divergent correlation predictions, undermining their interpretability. The current method eases this risk by maximizing entropy to select the least biased matrix under constraints (B-factors, symmetry, zero row sums, and fixed topology). These constraints significantly reduce the solution space compared to unconstrained fitting, enhancing the physical relevance of the results.

However, some indeterminacy persists due to the underdetermined nature of the system, highlighting the need for additional constraints.

Our approach provides a framework for systematically reducing indeterminacy by incorporating consequential constraints, drawing inspiration from statistical mechanics. For instance, experimental correlation data from NMR or single-molecule FRET could directly constrain off-diagonal elements, while side-chain interaction potentials or secondary structure-specific weights could ground adjustments in molecular properties. In statistical mechanics, similar strategies are well-established: the maximum entropy principle derives the canonical ensemble by constraining average energy, yielding the least biased distribution[37]; constrained Ising models use pairwise correlations to predict phase transitions[38]; and mean-field approximations in polymer physics, like Flory’s Gaussian chain model, reduce configurational indeterminacy with contact constraints[39]. General principles for entropy maximization subject to constraints is explained fully by Callen[40]. These precedents point to the power of constraints to transform underdetermined problems into well-defined solutions, positioning our method as a step toward a comprehensive solution for protein dynamics modeling. Future work should explore constraints like residue-specific interactions, long-range decay functions, or MD-derived correlations to further reduce indeterminacy, ensuring physically meaningful and unique Kirchhoff matrices.

The model’s limitations include its Gaussian fluctuation assumption, partially addressed by conditional mutual information but still potentially missing anharmonic motions, and its reliance on a fixed cutoff distance, which may miss long-range interactions in large systems. The computational cost of Monte Carlo optimization, while modest, could be a bottleneck for very large proteins, and B-factor accuracy depends on experimental data quality, potentially affected by crystal artifacts. Future work should explore non-Gaussian distributions, adaptive cutoffs, and machine learning to optimize residue assignments. Experimental validation of correlations and information flow, using NMR or single-molecule techniques, is essential to confirm predictions and address concerns about overfitting and indeterminacy. By addressing these challenges, the improved GNM provides a coarse-grained tool for studying protein dynamics and allostery.

## Supplementary Material 1

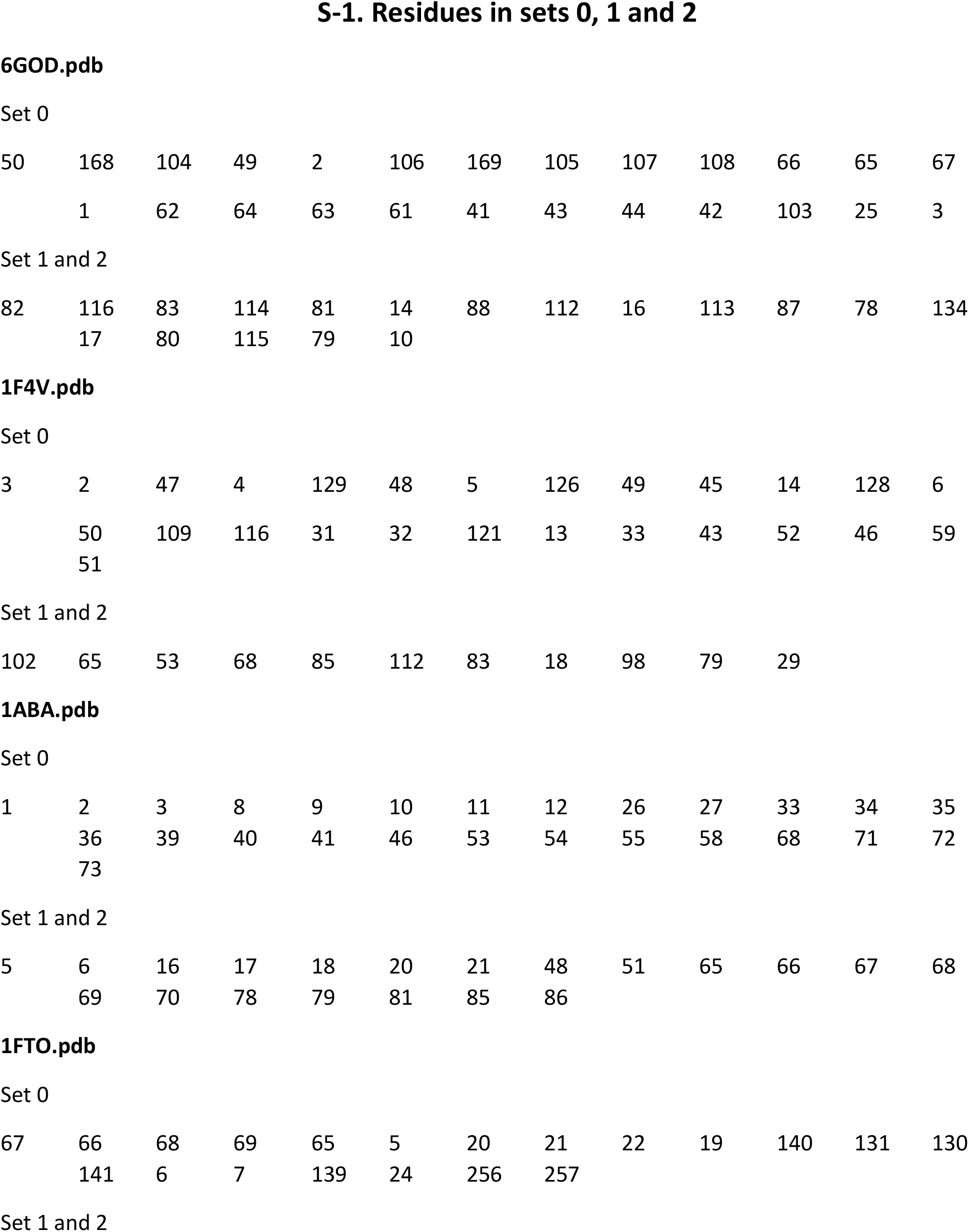
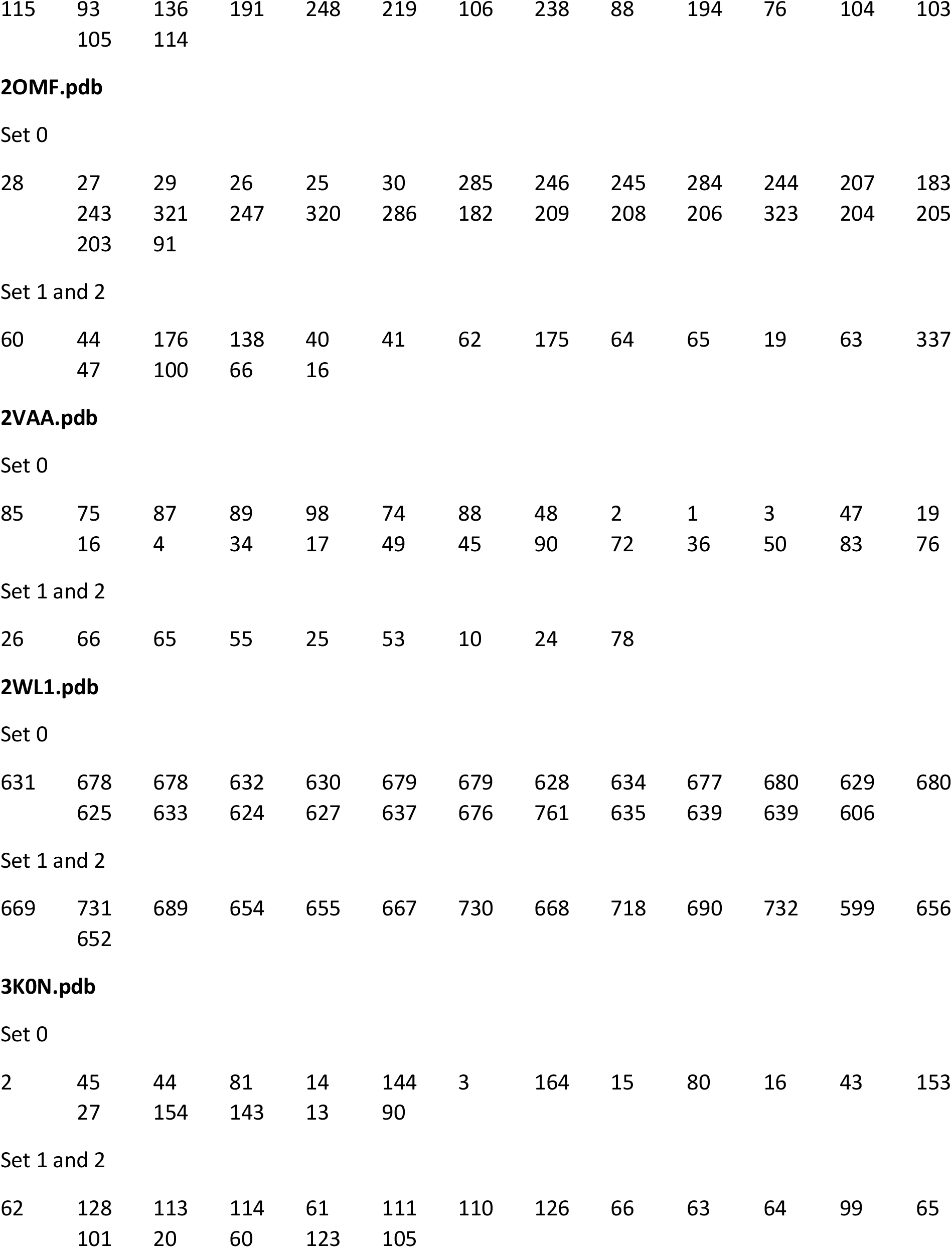

|  |  |  |  |  |  |  |  |  |  |  |  |  |
| --- | --- | --- | --- | --- | --- | --- | --- | --- | --- | --- | --- | --- |
| 50 | 168 | 104 | 49 | 2 | 106 | 169 | 105 | 107 | 108 | 66 | 65 | 67 |
|  | 1 | 62 | 64 | 63 | 61 | 41 | 43 | 44 | 42 | 103 | 25 | 3 |
| Set 1 and 2 |  |  |  |  |  |  |  |  |  |  |  |  |
| 82 | 116 | 83 | 114 | 81 | 14 | 88 | 112 | 16 | 113 | 87 | 78 | 134 |
|  | 17 | 80 | 115 | 79 | 10 |  |  |  |  |  |  |  |
| 3 | 2 | 47 | 4 | 129 | 48 | 5 | 126 | 49 | 45 | 14 | 128 | 6 |
|  | 50 | 109 | 116 | 31 | 32 | 121 | 13 | 33 | 43 | 52 | 46 | 59 |
|  | 51 |  |  |  |  |  |  |  |  |  |  |  |

Set 1 and 2
|  |  |  |  |  |  |  |  |  |  |  |
| --- | --- | --- | --- | --- | --- | --- | --- | --- | --- | --- |
| 102 | 65 | 53 | 68 | 85 | 112 | 83 | 18 | 98 | 79 | 29 |
| 1 | 2 | 3 | 8 | 9 | 10 | 11 | 12 | 26 | 27 | 33 | 34 | 35 |
|  | 36 | 39 | 40 | 41 | 46 | 53 | 54 | 55 | 58 | 68 | 71 | 72 |
|  | 73 |  |  |  |  |  |  |  |  |  |  |  |
| Set 1 and 2 |  |  |  |  |  |  |  |  |  |  |  |  |
| 5 | 6 | 16 | 17 | 18 | 20 | 21 | 48 | 51 | 65 | 66 | 67 | 68 |
|  | 69 | 70 | 78 | 79 | 81 | 85 | 86 |  |  |  |  |  |
| 67 | 66 | 68 | 69 | 65 | 5 | 20 | 21 | 22 | 19 | 140 | 131 | 130 |
|  | 141 | 6 | 7 | 139 | 24 | 256 | 257 |  |  |  |  |  |
| Set 1 and 2 |  |  |  |  |  |  |  |  |  |  |  |  |
| 115 | 93 | 136 | 191 | 248 | 219 | 106 | 238 | 88 | 194 | 76 | 104 | 103 |
|  | 105 | 114 |  |  |  |  |  |  |  |  |  |  |
| 28 | 27 | 29 | 26 | 25 | 30 | 285 | 246 | 245 | 284 | 244 | 207 | 183 |
|  | 243 | 321 | 247 | 320 | 286 | 182 | 209 | 208 | 206 | 323 | 204 | 205 |
|  | 203 | 91 |  |  |  |  |  |  |  |  |  |  |
| Set 1 and 2 |  |  |  |  |  |  |  |  |  |  |  |  |
| 60 | 44 | 176 | 138 | 40 | 41 | 62 | 175 | 64 | 65 | 19 | 63 | 337 |
|  | 47 | 100 | 66 | 16 |  |  |  |  |  |  |  |  |
| 85 | 75 | 87 | 89 | 98 | 74 | 88 | 48 | 2 | 1 | 3 | 47 | 19 |
|  | 16 | 4 | 34 | 17 | 49 | 45 | 90 | 72 | 36 | 50 | 83 | 76 |

Set 1 and 2
|  |  |  |  |  |  |  |  |  |
| --- | --- | --- | --- | --- | --- | --- | --- | --- |
| 26 | 66 | 65 | 55 | 25 | 53 | 10 | 24 | 78 |
| 631 | 678 | 678 | 632 | 630 | 679 | 679 | 628 | 634 | 677 | 680 | 629 | 680 |
|  | 625 | 633 | 624 | 627 | 637 | 676 | 761 | 635 | 639 | 639 | 606 |  |
| Set 1 and 2 |  |  |  |  |  |  |  |  |  |  |  |  |
| 669 | 731 | 689 | 654 | 655 | 667 | 730 | 668 | 718 | 690 | 732 | 599 | 656 |
|  | 652 |  |  |  |  |  |  |  |  |  |  |  |
| 2 | 45 | 44 | 81 | 14 | 144 | 3 | 164 | 15 | 80 | 16 | 43 | 153 |
|  | 27 | 154 | 143 | 13 | 90 |  |  |  |  |  |  |  |
| Set 1 and 2 |  |  |  |  |  |  |  |  |  |  |  |  |
| 62 | 128 | 113 | 114 | 61 | 111 | 110 | 126 | 66 | 63 | 64 | 99 | 65 |
|  | 101 | 20 | 60 | 123 | 105 |  |  |  |  |  |  |  |

## Notes

### Competing Interest Statement

The authors have declared no competing interest.

